# RNA-binding is the essential biological function of the *Drosophila* protein Brat

**DOI:** 10.64898/2026.02.27.708583

**Authors:** Robert P. Connacher, Yichao Hu, Richard Roden, Julia Toledo, Anna DesMarais, Michael O’Connor, Howard D. Lipshitz, Aaron C. Goldstrohm

## Abstract

Brain tumor (Brat) is a Drosophila TRIM-NHL protein required for embryogenesis and neural stem cell differentiation. Although structural and biochemical studies established that the Brat NHL domain specifically binds RNA, the in vivo requirement for this activity has not been directly tested. Here, we used structure-guided mutagenesis and genome engineering to determine whether RNA recognition is essential for Brat function during development. The direct interaction between Brat’s NHL domain and RNA containing Brat Binding Sites (BBS) can be abolished by alanine substitution of three separate residues on the NHL surface. We introduced these point mutations into the endogenous *brat* locus by CRISPR-mediated Scarless Gene Editing to generate three independent RNA-binding defective mutant (RBDmt) alleles. Complementation tests demonstrated that each allele behaves as a strong loss-of-function mutation: homozygotes and hemizygotes are inviable, and RBDmt alleles fail to complement classical *brat* null and hypomorphic alleles. Lethal phase analysis revealed death predominantly during late larval and pupal stages, consistent with known *brat* alleles. Consistent with the namesake *brat* phenotype, RBDmt larval brains exhibited widespread expression of neuroblast markers and a marked reduction of neuronal differentiation. In embryos, these alleles failed to complement female sterile *brat* alleles and recapitulated characteristic abdominal segmentation defects. Finally, RT-qPCR showed increased expression of endogenous Brat target mRNAs in mutant larvae, consistent with loss of Brat-mediated repression. Together, these results demonstrate that direct RNA binding is the essential molecular activity of Brat and that post-transcriptional regulation of Brat target mRNAs underlies its critical roles across development.

## Introduction

The *D. melanogaster* protein Brain tumor (Brat) mediates crucial developmental transitions during the fly life cycle including embryogenesis and differentiation of neural stem cells (reviewed in ^1^). Brat is a founding member of the TRIM-NHL family, consisting of an atypical, N-terminal tripartite motif (TRIM) domain, unstructured intervening sequence, and C-terminal NCL-1/HT2A/LIN-41 (NHL) domain (Figure 1(A)). Extensive biochemical and functional evidence demonstrates the NHL domain binds to specific RNA sequences (Figure 1(B)) ^2–5^. The 3′ untranslated regions (3′UTRs) of mRNAs that are both bound to ^3,4^ and regulated by Brat ^6^ are enriched in motifs bearing a 5′-UGUU core. These Brat Binding Sites (BBS) function as cis-regulatory elements, recruiting Brat to reduce mRNA stability and translation ^1^. Three-dimensional X-ray crystallography of Brat bound to BBS RNA revealed that residue F916 (Figure 1(C)) interacts with the central guanine of the motif via pi-stacking ^4^. Residue N933 (Figure 1(D)) hydrogen bonds with the adjacent uridine, while R875 (Figure 1(E)) forms ionic interactions with the adjacent backbone phosphates ^4^. Substitution of F916 or N933 to alanine eliminates the association of the NHL domain with the consensus sequence (5′-UUGUUGU_9_) or poly(U) RNA *in vitro* ^4^. Similarly, substitution of R875 to alanine strongly reduces binding to these sequences and fragments of known mRNA targets ^4,7^.

**Figure 1.**
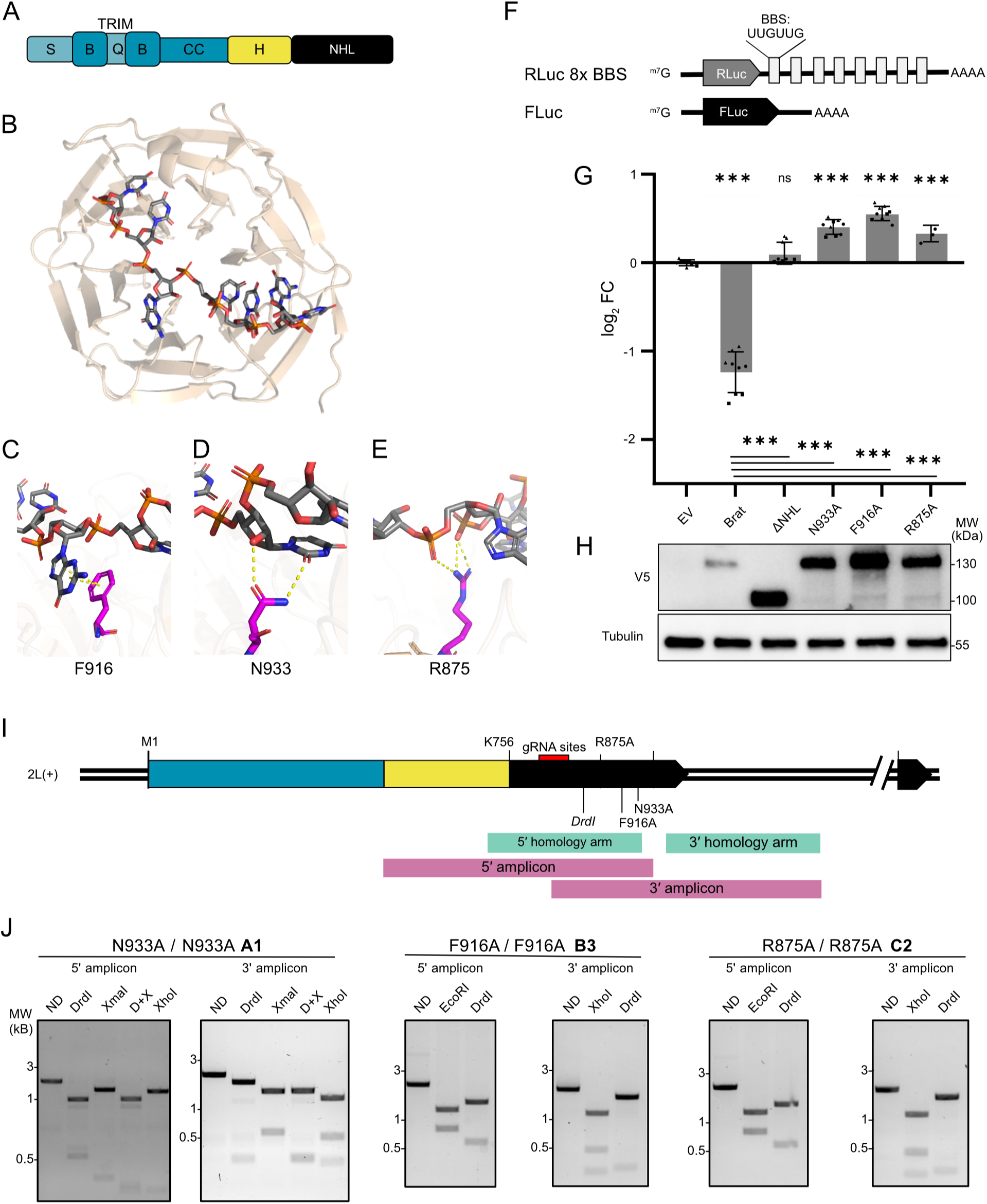
Design and validation of RNA-binding defective mutant *brat* alleles. A) Schematic diagram of Brat protein showing its domains. TRIM domain (blue) consists of the structure B-Box zinc fingers (B) and coiled-coil (CC) subdomains, as well as intrinsically disordered serine-rich (S) and glutamine-rich (Q) regions. The NHL domain (black) is connected to the coiled-coil domain via a histidine-ricoesh (H) region. B) X-ray crystal structure of the Brat NHL domain (ribbon structure, wheat) in complex with 5′-UUGUUGU RNA (ball and stick structure, grey and HNOS), PDB #5EX7. C) F916, D) N933, and E) R875 form non-covalent interactions with the RNA. Predicted hydrogen bonds (D, E) and pi-stacking interactions (C) shown as dashed lines. F) Renilla Luciferase (RLuc) reporter mRNA containing eight <u>B</u>rat <u>b</u>inding <u>s</u>ites (BBS) in the 3′UTR. Firefly Luciferase (FLuc) served as an internal control. G) Luciferase assay testing the ability of Brat or mutated versions to repress the reporter. The mean ± standard deviation of the log_2_ fold-change (FC), relative to Empty Vector (EV) controls, is plotted for different effectors. Significance of differences compared EV controls (top) or with wild-type Brat (bottom) determined via ANOVA with post-hoc Tukey–Kramer tests (n=9, *** p-value < 0.001, n.s.: not significant). H) Western blot of V5-tagged Brat proteins (bottom). Tubulin served as a loading control. Molecular weight markers are shown on the right in units of kDa. I) Diagram of the *brat* gene, denoting guide RNA (gRNA) sites, introduced mutations, and span of homology arms used for homologous recombination. Following Scarless Gene Editing, 5′ and 3′ regions covering the homology arm – genomic DNA junctions were amplified from the genomic DNA of homozygous larvae of the isogenic founder lines (bold) by PCR. J) Diagnostic restriction digests of PCR amplicons, followed by agarose gel electrophoresis, show the correct introduction of DrdI sites and XmaI sites (N933A only). Digestion with native XhoI or EcoRI sites were included as positive controls. ND (no digest), D+X (DrdI and XmaI). Molecular weight markers on the left are in kilobases.

Discovery of the RNA-binding activity of Brat prompted a reconsideration of its molecular and biological functions. In the initial model, developed from evidence in early embryos, Brat was proposed to serve as a translational repressor that is recruited to *hunchback* mRNA via protein-protein interactions with the RNA-binding proteins Pumilio and Nanos ^8,9^. Subsequently, the discovery that Brat binds to RNA independently ^4,7^ and the poor overlap with Pumilio targets ^3^ indicated that Brat directly selects its own BBS-containing target mRNAs. In addition, the scope of Brat’s molecular functions widened: Translational control predominates in the early embryo (^10,11^, and reviewed in ^12^), and at this stage Brat inhibits translation of *hunchback* mRNA without affecting mRNA stability ^8,9,13^. During the maternal-to-zygotic transition; however, Brat-associated mRNAs are destabilized ^3^. Brat similarly decreases the steady-state mRNA levels in other cell types ^5,6,14,15^.

In parallel, the role of Brat in neural stem cell differentiation has been continually evolving. In the larval brain, Brat ensures the proper differentiation of Type II neuroblasts (TIINBs) into neurons and glial cells in adults (reviewed in ^1,16,17^). When Brat is absent or reduced in TIINBs, daughter cells revert to stem-like states, and these highly proliferative cells develop into tumors ^5,18–22^. The namesake ‘brain tumor’ phenotype derives from the overgrown brain lobes of late third instar larvae ^23–32^.

Early evidence indicated the NHL domain was essential for Brat’s function, as weak phenotypes arise from non-synonymous codon changes in the NHL domain ^25^. As in embryos, Brat was believed to function as a translational repressor during TIINB asymmetric cell division, enriched in the appropriate daughter cell by interactions between the NHL domain and the scaffold protein Mira ^20,33^. Later, it was demonstrated that Brat represses mRNAs encoding transcription factors *myc*, *dpn*, and *zld* in the daughter cells of TIINBs ^5,34^. This repression is likely a direct effect, as the NHL domain binds to segments of their 3′UTRs *in vitro* and Brat represses reporter mRNAs bearing these 3′UTRs ^4^.

The prevailing model is that Brat’s NHL domain provides both the affinity and specificity for mRNAs encoding pro-stemness factors, and upon binding destabilizes these mRNAs by unknown means ^4–6,34–36^. Though accumulating evidence indicates that Brat controls mRNA fate in the germline, embryo, and larval brain, the precise role of its RNA-binding activity has not been directly tested in vivo. To date, the analysis of Brat in vivo has relied on either classical loss-of-function alleles, hypomorphic alleles, or depletion via RNAi.

In this study, we took a direct approach to determine the role of Brat RNA-binding activity in flies by introducing mutations into the endogenous *brat* locus, generating three separate RNA-binding defective mutant (RBDmt) alleles. We then tested the impact of these mutations on Brat’s functions and phenotypes. Our results show the RNA-binding activity of Brat is essential for viability. RBDmt alleles of *brat* mimic strong loss-of-function alleles and do not complement classical *brat* alleles. Furthermore, we demonstrate that RNA-binding is essential for the role of Brat in neural stem cell differentiation and in embryogenesis. In larval brains, RBDmt alleles produce tumors replete with neural stem cells at the expense of neurons, matching the reported *brat* phenotype. In embryos, RBDmt alleles fail to complement the *brat* female sterile alleles, which cause a characteristic loss of body segments. Finally, we show that mRNAs of Brat target mRNAs are derepressed by the RBDmt alleles in a manner consistent with loss of Brat repressive function. Taken together, these results demonstrate that post-transcriptional regulation of mRNAs is the primary biological function of Brat throughout development.

## Materials and Methods

### Fly genetics

Stocks containing the deficiency *Df(2L)Sd57* (Df, #101357) and *brat* alleles *brat{lacW}^k060^*^28^ (k06028, #114346) *brat*^18^ (#101379), *brat^ts1^* (#107544), *brat^fs1^*(#101352), and *brat^fs3^* (#101399) were obtained from the University of Kyoto Dept. Drosophila Genomics and Genetic Resources. Flies expressing *vasa>Cas9* (#55821 & #51324) or *nos>Cas9* (#78782), as well as *brat^1^* (#3988), were obtained from the Bloomington Drosophila Stock Center (BDSC). The *brat* locus of *brat^1^* and *brat*^18^ flies were amplified from hemizygous larvae using RC351 & RC357. Sanger sequencing identified the nonsense mutations G872* and Q507*, respectively. To confirm the reported lesions in female sterile *brat* alleles, the region of the NHL domain was amplified from several hemizygous *brat^fs1^/Df* and *brat^fs3^/Df* larvae using RC185 & RC369. Sanger sequencing with RC185 and RC269 verified the missense mutations G774D and H802L, respectively (Figure S3(B)). Wild-type flies were *w^11^*^18^ or *y^-^,w^-^*. Flies containing the PiggyBac transposase (BDSC #8283) were kindly provided by Tom Hayes.

### Molecular cloning

All plasmids, oligonucleotides, and synthesized DNA fragments used in this study are listed in (Additional File 1). RNA-binding mutations were introduced into the pIZ Brat via QuickChange Site-Directed Mutagenesis (Agilent) using oligos RC173/RC174 (N933A), RC419/RC420 (F916A), or RC417/RC418 (R875A). pIZ ΔNHL (Δ756-1040) was derived from pIZ Brat via inverse PCR, using oligos RC184 & RC188. Guide RNAs targeting the NHL domain of Brat were identified via flyCRISPR’s Optimal Target Finder tool for BDSC #51324 ^37^. The guide RNA sequences were cloned into pCFD4-U6-1_U6:3tandemgRNAs (Addgene #49411) via Gibson assembly. Specifically, the large BbsI fragment was fused to a fragment containing both guide RNAs and the dU6-1 – 3 intervening sequence (amplified from the small BbsI fragment via RC_gRNA_F & RC349).

The homology-directed repair template was generated from pScarlessHD-DsRed-w+ (Addgene #80801). Gibson assembly was used to combine five fragments: 5′ homology arm (amplified from BDSC #55821 genomic DNA with RC286 & RC287), edited region, DsRed marker (amplified from pScarlessHD with RC290 & RC291), 3′ homology arm (amplified from genomic DNA with RC288 & RC311), and the plasmid backbone (amplified from pScarlessHD with RC310 & RC293). Specifically, the DsRed marker contained the promoters, DsRed, and SV40 poly(A) signal from pScarless, flanked by PiggyBac inverted repeats.

For F916A and R875A, the edited region was amplified from synthetic DNA fragments G_F916A and G_R875A with RC294 & RC295. For N933A, the edited region was amplified with RC294 & RC296. These fragments contained the mutation of interest as well as a 5′-TTAA, disrupted guide RNA sequence, and restriction site(s) introduced via synonymous re-coding. As necessary, a non-synonymous codon present in the 5′ homology arm was corrected via QuickChange mutagenesis (Agilent) using RC347 & RC348.

To generate Brat transgenes, the Brat coding sequence with C-terminal V5 and His6 epitope tags was amplified from pIZ Brat or pIZ Brat[N933A] ^6^ via RC800 & RC801, and inserted into the KpnI & XbaI sites of pUAS-1.0 ^38^ via restriction cloning. Transgenic flies were generated via PhiC31 integration of these transgenic vectors into embryos bearing attP2 sites (BDSC #8622), by TheBestGene.

### Fly CRISPR

CRISPR was conducted via Scarless Gene Editing ^39^. HDR template and gRNA plasmids were injected into *nos-Cas9* embryos (#TH00787.N) by TheBestGene. Founders were mated with *y^-^w^-^* flies, and progeny were screened for expression of DsRed and lack of mini-white in eyes. Isogenic lines were maintained from appropriate founders. For all lines, multiple founder lines were maintained. The junction of genomic DNA and homology arm was verified by amplifying the region of interest with RC185 / MJ01 or RC401 / 402. Once confirmed, the dsRed cassette was mobilized using PiggyBac transposase, and isogenic lines once again developed from individual founders. Extensive confirmation of appropriate edits involved amplifying the genomic DNA of homozygous or hemizygous larvae via RC368 / 406 or RC185 / 369, followed by restriction digests and Sanger Sequencing. All identified single nucleotide polymorphisms are listed in (Additional File 1). Notably, a non-synonymous mutation of G643S was observed in several lines; but this occurs often in natural *Drosophila* populations ^40,41^ and likely derived from the *nos-Cas9* stock.

### Luciferase assays

Luciferase assays were conducted as described ^6^. Assays were conducted with three biological replicates (each transfected well), and three experimental repeats (independent experiments performed on separate days). Renilla and Firefly luminescence in each well was quantified via a Dual-Glo Luciferase Assay system (Promega) and a Glomax Discover (Promega) instrument. For each sample, the Renilla luminescence signal was normalized to that of the internal control Firefly. Then, for each effector protein, the ratio was divided by the mean ratio of the empty vector negative control to calculate the fold change caused by the effector protein. The resulting values (log_2_ transformed) were fit to a linear model with mixed effects, in order to preserve variation from both biological and experimental sources. Statistical differences between groups were determined via ANOVA with post-hoc Tukey-Kramer tests. The datasets and outputs of these tests are presented in (Additional File 1). Expression of the transfected proteins was confirmed via western blotting, as described ^6^.

### Fly viability assays

The complete results of all fly assays are presented in (Additional File 1). To test the viability of flies with different combinations of *brat* alleles, 5-10 virgin females were mated with an equal number of males in cornmeal-agar vials. In all cases, the *CyO,Star^1^* derivative of the common *CyO* balancer was used, as it incorporates a dominant *Star^1^* eye phenotype in addition to *Cy* wings. Flies were moved to new vials every two days to prevent crowding, for three to four total passages. After 10 days at 25°C, progeny adults were cleared daily until 20 days after egg laying, and the total *Cy* and non-*Cy* flies counted. Unless otherwise noted, experiments were conducted with triplicate crosses and counts of total progeny combined. Accounting for the embryonic lethality of balancer homozygotes, the expected proportion of countable progeny with a non-lethal, somatic allele combination in flies is 0.3 following Mendelian inheritance ^42^. The difference between the observed counts and the expected counts for the same number of progeny was determined via a Chi-Squared Test. P-values were corrected via the Bonferroni method to account for multiple testing. Stacked bar graphs were generated in R using the *ggplot2* package. The full counts, original and adjusted p-values, and number of replicate assays are detained in Additional File 1

To investigate the lethal phase of *brat* mutants, flies of the appropriate genotypes were mated in cages over apple juice agar plates, with eggs laid in approximately 24 hour windows. The resulting embryos were allowed to age an additional 24 hours until larvae hatched. In these assays, the *CyO,GFP* derivative of the *CyO* balancer was used, so genotypes of these larvae could be deduced by a strong band of green fluorescent protein (GFP) expression in the abdomen of heterozygotes. Twenty GFP^-^ or GFP^+^ larvae were placed in cornmeal-agar vials and incubated at 25°C. Immobile pupae were marked until 10 days after egg laying, and adult flies were removed/counted daily until 20 days after egg laying. Care was taken to notice any inappropriate *Cy* flies in the non-*Cy* vials, and if found, the total number of initial larvae adjusted accordingly. Experiments were conducted in triplicate from the same initial cross (n=3); however conducting similar experiments with triplicate crosses (n=3x3) produced similar results (Figure S3). Clustered bar graphs were generated in R using the *ggplo2* package. A Welch Two Sample t-test was used to assess the significance of differences in the percentage of larvae that survived to pupal or adult stages, comparing *brat* mutants and their heterozygous siblings.

To test haploinsufficiency of *brat* alleles, flies heterozygous for *brat* alleles and the *CyO,GFP* balancer chromosome were mated to wild-type (*w^11^*^18^) flies. Flies were moved to new vials daily, for a total of three passages, with each functioning as a technical replicate. Crosses were conducted in triplicate. The fraction of non-*Cy* adult progeny that eclosed between 10 and 20 days after egg laying was determined for each vial. A one-way ANOVA determined the differences between these fractions were not significant (p-value = 0.1573).

To assess viability following transgenic over-expression of *brat* or derivatives, flies homozygous for the *daughterless>Gal4* driver were crossed to flies heterozygous for transgenes and the chromosome III balancer *TM3, Sb*. The resulting non-*Sb* adult progeny ubiquitously over-express Brat (or N933A). The fraction of *non-Sb* progeny was compared between these, and to similar crosses with a *TM3, Sb* donor that lacked transgenes. These crosses were conducted in triplicate, with flies moved to new vials daily for a total of three passages. The total adult progeny of these technical replicates were combined, for each experimental replicate.

### Embryonic cuticle preps

Females of the appropriate genotype laid eggs on apple juice agar plates in approximately 24 hour windows. The resulting embryos were allowed to age an additional 24 hours until late embryonic / early larval stages, then collected and dechorionated in 5 mL of 2.5% sodium hypochlorite for two minutes, followed by two 10 milliliter washes with water. Dechorionation and washing utilized a vacuum filter with a nylon mesh membrane. Embryos were incubated, on microscope slides, in a 1:1 mixture of lactic acid and Hoyer’s medium at 65°C overnight ^43,44^. The resulting cuticles were imaged in dark-field to visualize abdominal denticle bands.

### Immunohistochemistry and microscopy

Brains of wandering third instar larvae were dissected in cold phosphate buffered saline (PBS) and collected into 1.5mL centrifuge tubes (all experiments were conducted at room temperature if not specifically stated). Samples were fixed for 20 minutes in Fixing Buffer (4% paraformaldehyde, 0.1% Triton-X100, PBS). Samples were washed three times for 10 min each in Wash Buffer (0.1% Triton-X100, PBS), and then blocked for one hour in BSA Blocking Buffer (1% bovine serum albumin, 0.1% Triton-X100, PBS). Samples were incubated with primary antibody solution (primary antibodies diluted in BSA Blocking Buffer) overnight at 4°C. Primary antibodies used included rat anti-Elav (1:50, Developmental Studies Hybridoma Bank), rat anti-Brat (1:250 ^20^), rabbit anti-Brat (1:50 ^9^), rabbit anti-Mira (1:200 ^21^), and rat anti-Dpn (1:100, Abcam #ab195173). Samples were washed three times for 10 min each in PBST, then incubated in Secondary Antibody Solutions (Alexa Flour secondary antibodies diluted 1:300 in BSA blocking buffer) for two hours. Samples were washed three times for 10 min each with PBST, then transferred to antifade mounting medium (2.5% DABCO in 70% glycerol) for at least four hours. Samples were mounted in an antifade mounting medium immediately before imaging. Images were collected using a Leica DMi8 TCS SP8 confocal microscope and captured with the LAS X software.

### Reverse transcription - quantitative PCR (RT-qPCR)

Crosses were conducted in mating cages over apple juice agar plates, using flies with a *brat* allele balanced with *CyO,GFP* and another *brat* allele balanced with *CyO,Tb-RFP*. The latter is a derivative of the *CyO* balancer that expresses the *Tb^1^* fragment fused to red fluorescent protein (RFP). The resulting eggs were collected in six-hour windows approximately 24 hours later. First instar larva were placed in vented 6 cm dishes containing cornmeal-agar media. Five days after egg laying, larvae were floated in 20% sucrose, transferred to a dissecting dish, and washed in three wells of cold PBS. Following a final transfer to PBS, flies were sorted by expression of GFP, RFP, and tubby morphology (details in (Additional File 1)). For each genotype, RNA was isolated from pairs of larvae, with four pairs considered as four biological replicates. The same procedure was followed for wild-type larvae (*w^11^*^18^). RNA was isolated, cDNA transcribed, and RT-qPCR conducted following the procedures previously optimized to measure *VhaPPA1-1*, *Vha100-2*, *VhaM8.9*, and *Treh* mRNAs from larvae ^6^. Significance of differences in the levels of these mRNAs between genotypes was determined via ANOVA (anova() of *stats* package) and post-hoc Tukey-Kramer tests (ghlt() of *multcomp* package) in R. The underlying data and complete results of these comparisons are presented in (Additional File 1). The results and procedures are reported according to the MIQE guidelines ^45^, detailed in (Additional File 1).

## Results

### Generating RNA-binding defective alleles of brat

We sought to confirm that the RNA-binding residues identified biochemically and structurally were essential for the ability of Brat to repress a target mRNA. For this task, we employed a previously established dual luciferase reporter assay in cultured Drosophila cells that measures the repressive activity of Brat ^6^. A Renilla luciferase (RLuc) coding sequence was appended with a minimal 3′UTR containing eight Brat binding motifs (5′-UUGUUG) to generate Rluc 8xBBS (Figure 1(F)). As an internal control, a Firefly luciferase (FLuc) plasmid was co-transfected with the Renilla reporter plasmid into D.mel-2 cells. Decrease in the normalized RLuc luminescence signal, calculated relative to the empty vector (EV) control, measured the relative repression of the reporter by Brat. In these assays, the cells were co-transfected with plasmids expressing either wild-type Brat, Brat with the NHL domain deleted (ΔNHL), or Brat with RNA-binding mutations N933A, F916A, or R875A. Wild-type Brat effectively repressed the 8xBBS reporter (Figure 1(G)). In contrast, the mutant Brat proteins did not repress, similar to ΔNHL, despite being expressed at higher levels than wild-type Brat (Figure 1(H)). These observations are consistent with previous results, wherein Brat^R875A^ and Brat^F916A^ were shown to be less effective than wild-type Brat at repressing a reporter bearing the *hunchback* 3′UTR ^7^. Similar observations were made for Brat^N933A^, Brat^R875A^, and Brat^F916A^ with reporters bearing the 3′UTRs of the Brat target genes *Klumpfuss* or *Knirps* ^4^. These results highlight two crucial observations: First, mRNA repression by Brat depends on the NHL residues N933, F916, and R875 binding specific BBS mRNA motifs. Second, these RNA-recognition residues are non-redundant, as mutation of each amino acid alone prevents repression by Brat.

To determine the role of Brat’s RNA-binding activity during development. We introduced alanine substitutions of each of the three RNA-binding residues (N933, F916, R875) into the endogenous *brat* gene via scarless CRISPR gene editing. All isoforms of *brat* annotated in the r6.65 genome assembly share a common coding sequence except isoform E, which has a short C-terminal extension beginning at V1035. The position of the RNA-binding residues ensures that all isoforms of Brat contain the alanine substitutions. Appropriate editing of one founder for each allele (*brat^N933A^* A1, *brat^F916A^* B3, and *brat^R875A^* C2) was verified by PCR and Sanger sequencing (Figure S1, Additional File 1). Specifically, amplicons were designed to cover the genomic DNA-homology arm junctions, the entirety of the homology arms, and the edited region (Figure 1(I)). As a marker of editing, a DrdI site was introduced using synonymous codons. Digestion of the 5′ and 3′ amplicons with DrdI, or obligate cutter EcoRI (5′) or XhoI (3) produced the expected banding patterns (Figure 1(J)). Additionally, the N933A edit introduced an XmaI site - which similarly produced the expected band patterns upon digest. The combination of sequencing and restriction digests confirmed that we had successfully engineered flies with the three different RNA-binding defective alleles of Brat.

### RNA-binding by Brat is essential for viability

To test the broad importance of RNA-binding, we assessed the impact of the RNA-binding defective *brat* alleles on viability via complementation assays. In these experiments, the parents are heterozygous for *brat* alleles and a balancer chromosome (*CyO,Star^1^*). These balancer chromosomes carry a wild-type copy of *brat,* a dominant and readily scorable *Duox^Cy^* and *Star^1^* marker, suppress meiotic recombination, and contain embryonic lethal, recessive alleles ^46^. In a test cross of two balanced *brat* alleles (Figure 2(A)), two lethal recessive alleles that cannot complement are expected to produce 100% *Cy Star^1^* (heterozygous) offspring. Alternative, non-lethal, complementary allelic combinations are expected to produce 33% non-*Cy Star^1^* (homozygous) offspring. For the observed ratio and sample size, the chi-squared goodness of fit test was used to determine whether the percentage of homozygous progeny was significantly different from the expected non-lethal allelic combination percentage. Such assays were originally used to identify and assess *brat* alleles ^23,24^.

**Figure 2.**
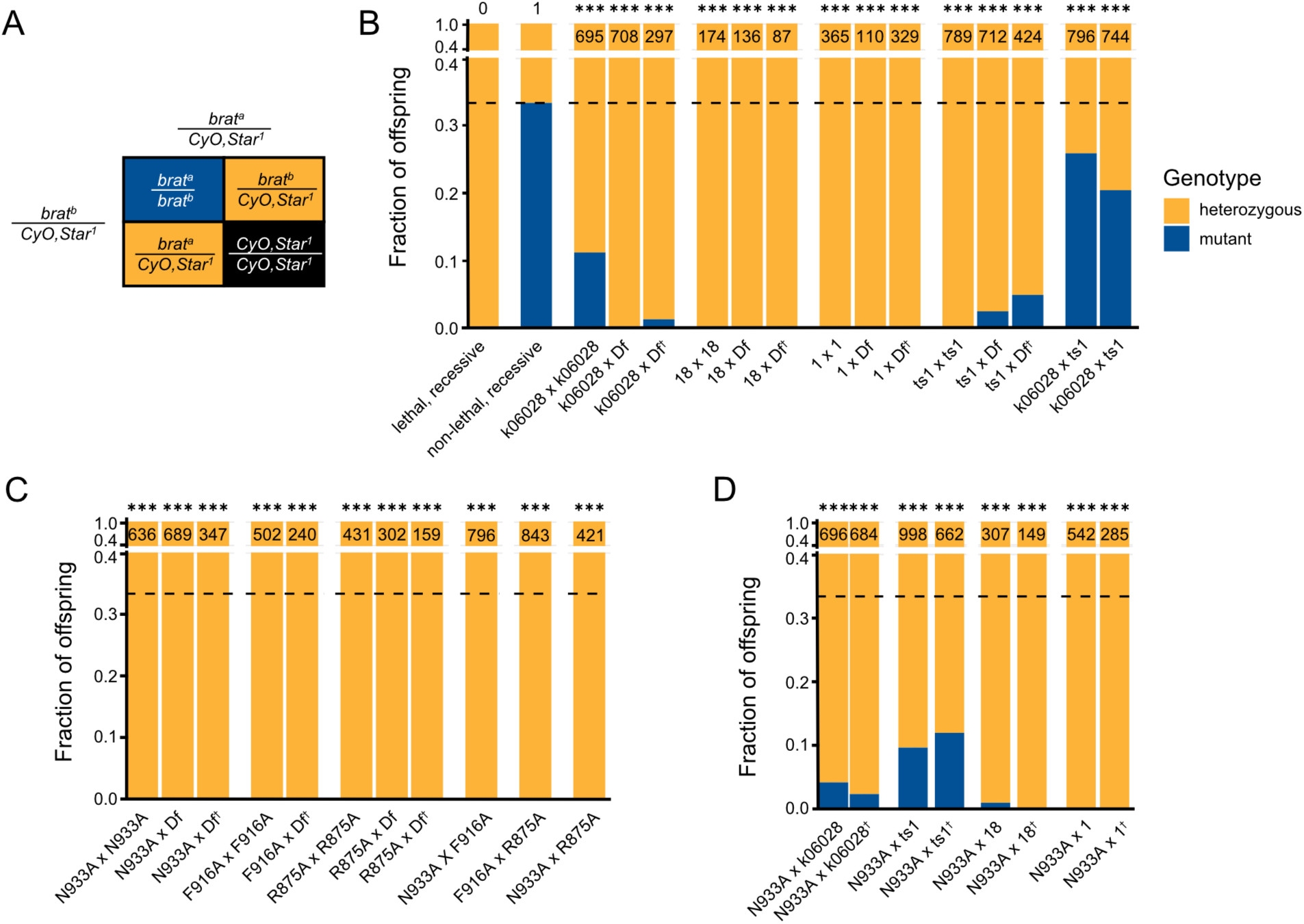
*Brat* RBDmt alleles do not complement known loss of function alleles. A) Punnett square displaying the expected outcomes of a monohybrid cross using balancer chromosomes. In these crosses, heterozygous offspring (yellow) are marked with curly wings and the *Star^1^* eye phenotype. B) Stacked column graphs showing percentage of heterozygous (yellow) or mutant (blue) offspring of the indicated cross. Total offspring counted from all replicate crosses shown above. The expected outcomes of a lethal, recessive and non-lethal allelic combination are noted. The observed fraction was compared to the expected fraction from a non-lethal combination (0.33, dashed line) by a χ^2^ Test with Bonferroni-corrected p-values for multiple comparisons (*** p^adj^ < 0.001). When relevant, crosses were also conducted with parental genotypes switched (†). Similar plots showing combinations of RBDmt alleles (C) and *N933A* with known *brat* alleles (D).

We first validated the complementation assay by assessing the viability of known *brat* alleles (Figure 2(B), Figure S2(A)). As expected, flies with known *brat* loss-of-function alleles similarly produced few homozygous offspring (Figure 2(B)). Alleles encoding premature termination codons that truncate Brat protein, such as *brat^1^* or *brat*^18^, were 100% lethal when homozygous. The same result was obtained when alleles were hemizygous with a chromosome deletion removing the *brat* coding sequence (*Df(2L)Sd57*, or *Df)*. The *k06028* allele contains a P-element insertion into the 5’UTR ^25^. This weaker, hypomorphic allele produced homozygous and hemizygous escaper adults, but at significantly lower rates than expected for a non-lethal combination. Finally, survival was greatest when a weaker, temperature-sensitive allele (*brat^ts1^*) was hemizygous; or complemented with *k06028,* producing nearly non-lethal rates of 20-26% trans-heterozygotes. Therefore, these complementation assays quantitatively assess the lethality of *brat* alleles.

We then tested the viability of RNA-binding defective *brat* alleles in both homozygous and hemizygous combinations. The RNA-binding mutants *N933A*, *F916A*, and *R875A* were all 100% lethal as homozygotes and when complemented by a deficiency (Figure 2(C)). Similar results were obtained for additional founder alleles of *N933A*, *F916A*, and *R875A* (File S2(B-D)). Combinations of these alleles were also lethal (Figure 2(C)). To ensure that the defect in viability was due to a lack of functional *brat*, *N933A* was crossed to several known *brat* alleles (Fig 2(D)). In all cases, *N933A* acted as a strong loss-of-function allele, with percentages of homozygous adults like those obtained when the alleles were complemented with a deficiency. These results indicate that RNA-binding is essential for the function of Brat, and point mutants that interrupt RNA-binding are indistinguishable from protein-null alleles.

### RNA-binding by Brat is required in larval and pupal stages

The *Drosophila* life cycle (Figure 3(A)) begins with embryos, which hatch as larvae, develop through three instars (L1-L3), and subsequently pupariate ^46^. These pupae undergo metamorphosis, finally eclosing as adults. While embryonic phenotypes of *brat* mutants have been described ^3,8,47^, these phenotypes are complicated by lingering maternally-deposited *brat* mRNA and protein. The first identified phenotype caused by loss of zygotically produced Brat alleles occurs in the neural stem cells of late L3 larvae ^20^. To identify the stages in which *brat* RBDmt alleles die, we conducted lethal phase analysis. In these experiments, test crosses used flies heterozygous for *brat* alleles and a balancer chromosome marked with green fluorescent protein (GFP) (Figure 3(B)). The genotype of the resulting progeny was visually differentiated by the presence or absence of GFP, which is expressed throughout L1 larvae. An equivalent number of *brat* mutant (*brat^a^ / brat^b^*) first instar larvae - or their heterozygous siblings - were housed separately in vials and allowed to proceed through development. As pupae are immobile and can be non-invasively scored, and adults easily separated, we recorded the percentage of larvae that completed these key developmental transitions. Additionally, separating larvae and maintaining equivalent numbers in vials reduces larval crowding, which negatively affects survival through adult stages ^48^. Therefore, lethal phase analysis both provides information on timing, and serves as a parallel approach for testing the effect of *brat* alleles on viability independent of crowding.

**Figure 3.**
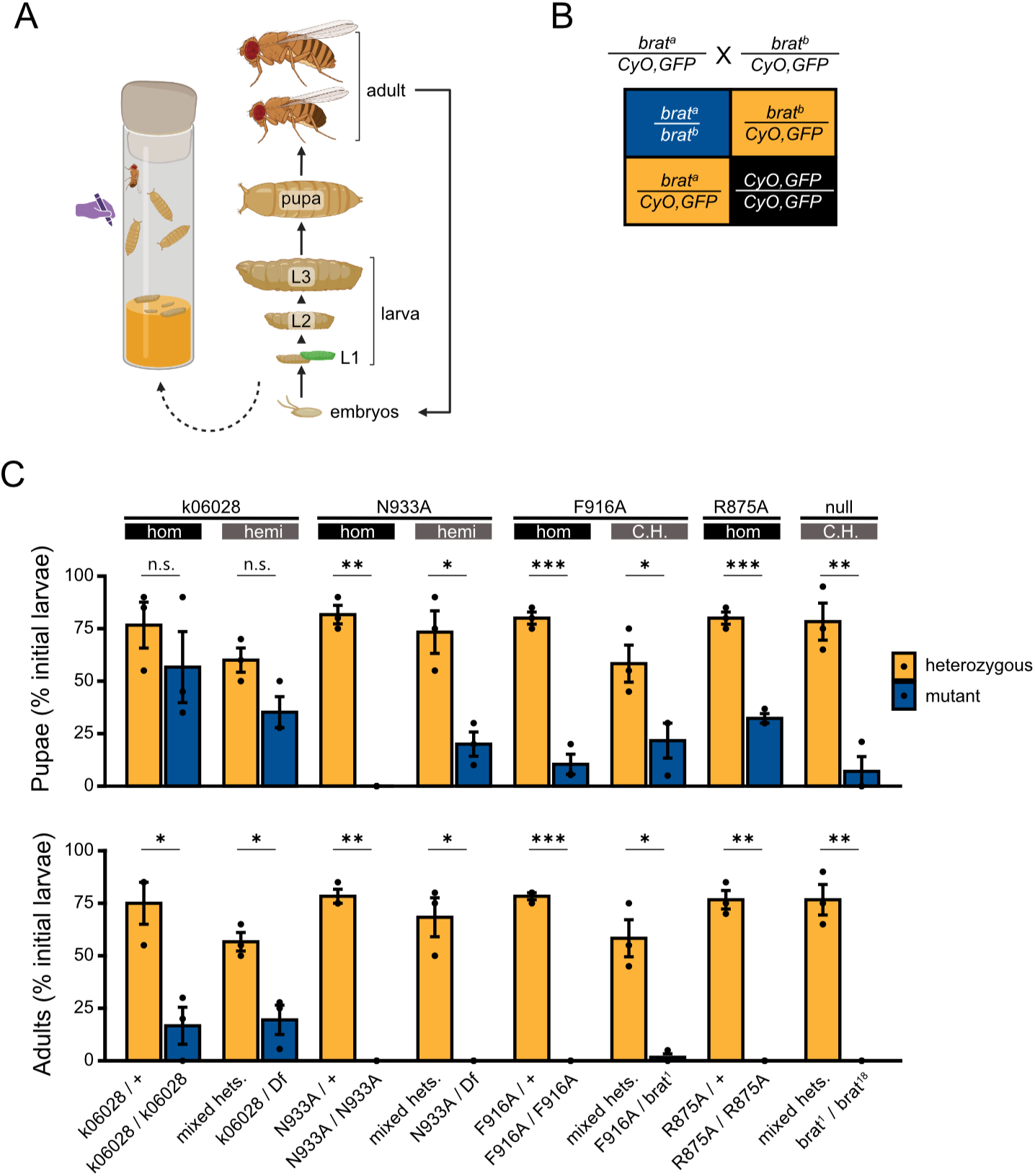
The lethal phase of RBDmt *brat* alleles occurs in late larval and pupal stages. A) Diagram of the *Drosophila* life cycle, noting when animals were sorted and counted. Created in BioRender. Connacher, R. (2026) https://BioRender.com/6dl60ofB B) Punnett square displaying the expected outcomes of a monohybrid cross using balancer chromosomes. In these crosses, heterozygous larvae (yellow) are marked with green fluorescent protein (GFP). C) Percentages of initial larvae that survive to pupal stages (top) and adulthood (bottom), as a percentage of initial larva. The offspring of the test crosses may be homozygous (hom), hemizygous (hemi), or compound heterozygotes (C.H.), noted above each comparison.

As a proof of principle, we first conducted assays with known *brat* alleles. Generally, larvae with *brat* allelic combinations either die in a stalled third instar state or non-eclosing pupae (Figure 3(C)). For example, larvae homozygous and hemizygous for *k06028* and hemizygous pupariated and eclosed at lower rates than their heterozygous siblings. This is consistent with observations that homozygous *k06028* animals develop brain tumors in the third larval instar ^25^, and the existence of rare adult escapers^49^. The phenotype was more extreme in a combination of null alleles (*brat^1^/*^18^), in which few larvae survived to pupariation. These assays used technical replicates from a single parental cross, and the observations were reproducible across repeated experiments (Figure S3(A-B)), and confirmed the essential role of *brat* in the late larval stages.

Complementation assays were then performed with the RNA-binding mutant alleles *N933A*, *F916A*, and *R875A* (Figure 2(C)). In all cases, few homozygous or hemizygous larvae survived to the pupal stage, and none survived to adulthood. It is notable that *N933A* homozygotes fared worse than hemizygotes, indicating an additional lethal, recessive allele on chromosome II. As this could be a confounding factor, further complementation assays used different combinations of *brat* alleles. Similar lethality was observed with the compound heterozygote *brat^F9^*^16^*/^1^*, indicating the phenotype is genuinely due to defects in the *brat* gene. Taken together, these results demonstrate that RNA-binding is essential for the function of Brat in late third instar larva and pupae.

### RNA-binding by Brat is required for neural stem cell differentiation

We next investigated the role of RNA-binding in the namesake *brat* phenotype. The central brain of third instar larva consists of a ventral nerve cord (VNC) and two central brain lobes that house neural stem cells (Figure 4(A)). The tumors of *brat* mutants originate in Type II neuroblasts (TIINBs) ^22^. While wild-type larva have approximately eight TIINBs per lobe ^22^, loss or reduction of Brat in these cells cause an overgrowth of neuroblast-like cells ^5,18^, which eventually leads to tumors ^19–21^. We investigated whether RNA-binding is essential for Brat’s function in neuroblasts by performing immunofluorescence microscopy (IF) on the brain lobes of RBDmt and heterozygous larvae. As a marker, the Dpn protein is expressed in larval neural stem cells (and in secondary neuroblasts), but not immediate progeny ^50,51^. In the brain lobes of *F916A* heterozygotes, Dpn expression was sparse (Figure 4(B), left panel). In contrast, Dpn expression was widespread in the (markedly larger) lobes of *F916A* homozygotes (Figure 4(B), right panel). Thus, loss of the RNA-binding function of Brat alone produces supernumerary neuroblasts. This is consistent with previous observations when *brat* is altogether absent ^20^, and demonstrates the importance of RNA-binding in Brat’s role as a tumor suppressor.

**Figure 4.**
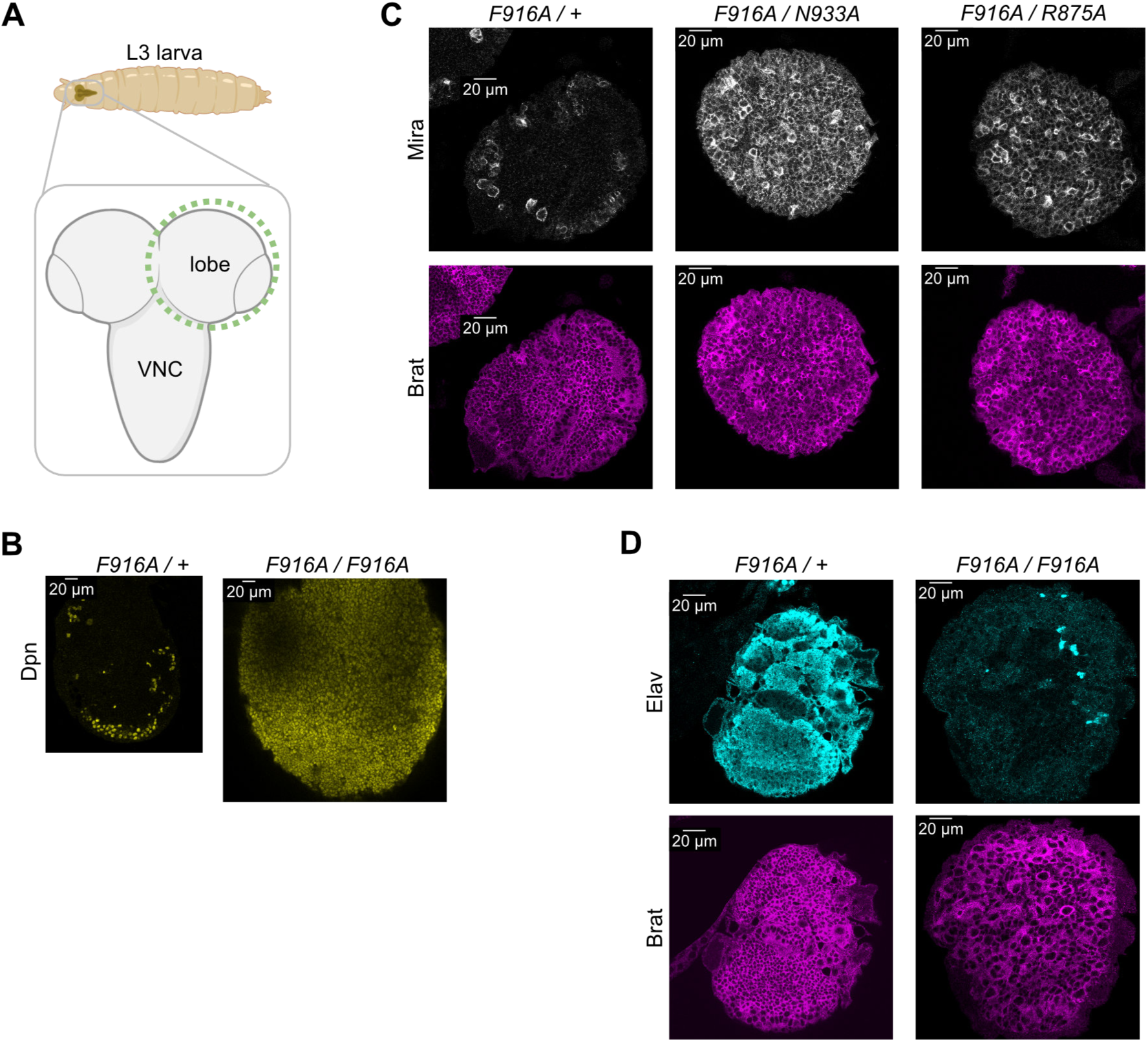
RBDmt *brat* alleles develop brain tumors. A) Diagram of the central nervous system (inset) within third instar larva. Type II neuroblasts are contained in the two brain lobes (dashed, green outline), but not the ventral nerve cord (VNC). Created in BioRender. Connacher, R. (2026) https://BioRender.com/am44do4 B-D) Immunofluorescence microscopy showing Dpn (B), Mira and Brat (C), or Elav and Brat (D) expression in a single brain lobe. Genotype of the larva is labeled above each panel.

Because Brat directly binds to and represses Dpn expression ^5,34^, we opted to verify this observation with additional neuroblast markers. Mira is predominantly expressed in the neural stem cells of the larval brain ^52^, where it is crucial for the asymmetric division that initiates neuroblast differentiation ^53,54^. The proportion of Mira^+^ cells per brain lobe is substantially higher in protein null *brat* mutants ^20^. We found that, while the brain lobes of heterozygous *F916A/+* larvae had few Mira+ cells, these cells were widespread in *N933A / F916A* and *R875A / F916A* larvae (Figure 4(C)). As *brat* tumors arise from both a block in neuroblast differentiation and overproliferation of these cells (reviewed in ^16^), we additionally probed for markers of neuroblast progeny. Elav is uniquely expressed in neurons but not glia or neuroblasts ^55^, and Elav^+^ cells are reduced in the brains of *brat* mutants ^20^. We observed that Elav+ cells were widespread in F916A heterozygotes and nearly absent in homozygotes (Figure 4(D)). To ensure that the phenotype was due to Brat dysfunction rather than reduced protein expression, we performed immunofluorescence detection of Brat protein in parallel. Despite a specific function in TIINBs ^22^, Brat is expressed broadly in central brain lobes (Figures 4(C-D)), as observed by ^21^. Crucially, Brat was expressed throughout both heterozygous and mutant brains, indicating the RNA-binding mutations do not adversely affect expression of Brat. These results demonstrate that the brain tumor phenotype of *brat* null mutants, driven by neuroblast overproliferation at the cost of differentiated progeny, is recapitulated by mutations that block Brat’s RNA-binding activity. We conclude that the RNA-binding function of Brat is essential for its role as a tumor suppressor and in promotion of neural stem cell differentiation.

### Brat RNA-binding activity is required in the early embryo

We next investigated the role of RNA-binding in additional, established *brat* phenotypes. Maternally provided Brat contributes to the regulation of maternally-deposited *hunchback* mRNA in early embryos, helping to establish body patterning in conjunction with the Pumilio and Nanos RBPs (reviewed in ^56^). Brat binds to specific sites in the *hunchback* mRNA ^7^ and misregulation of *hunchback* produces a defect in body segmentation ^8,9^. Therefore, we investigated whether the RNA-binding defective *brat* alleles produced a similar phenotype.

As a technical note, the lethal phase of zygotic *brat* mutants occurs during larval and pupal development (Figure 3). However, the ‘female-sterile’ class of alleles consist of weaker missense mutations that produce phenotypes in the ovaries of adult females and offspring ^3,8,9^. We initially tested whether the female-sterile allele *brat^fs1^* ^8^ could complement the RBDmt alleles, as in Figure 2. Flies with combinations of *brat^fs1^* and either *brat^k060^*^28^ or the temperature-sensitive *brat^ts1^* allele largely survive into adulthood; while flies hemizygous for *brat^fs1^* or in combination with RBDmt alleles (*brat^fs1^ / brat^N933A^*) are subviable (Figure 5(A)). As with previous compound heterozygotes (Figure 2(D)), *N933A* functioned like a strong loss-of-function allele of *brat*. Furthermore, the substantial fraction of *brat^fs1^/^N933A^* progeny obtained allowed us to investigate the phenotypes in progeny embryos.

**Figure 5.**
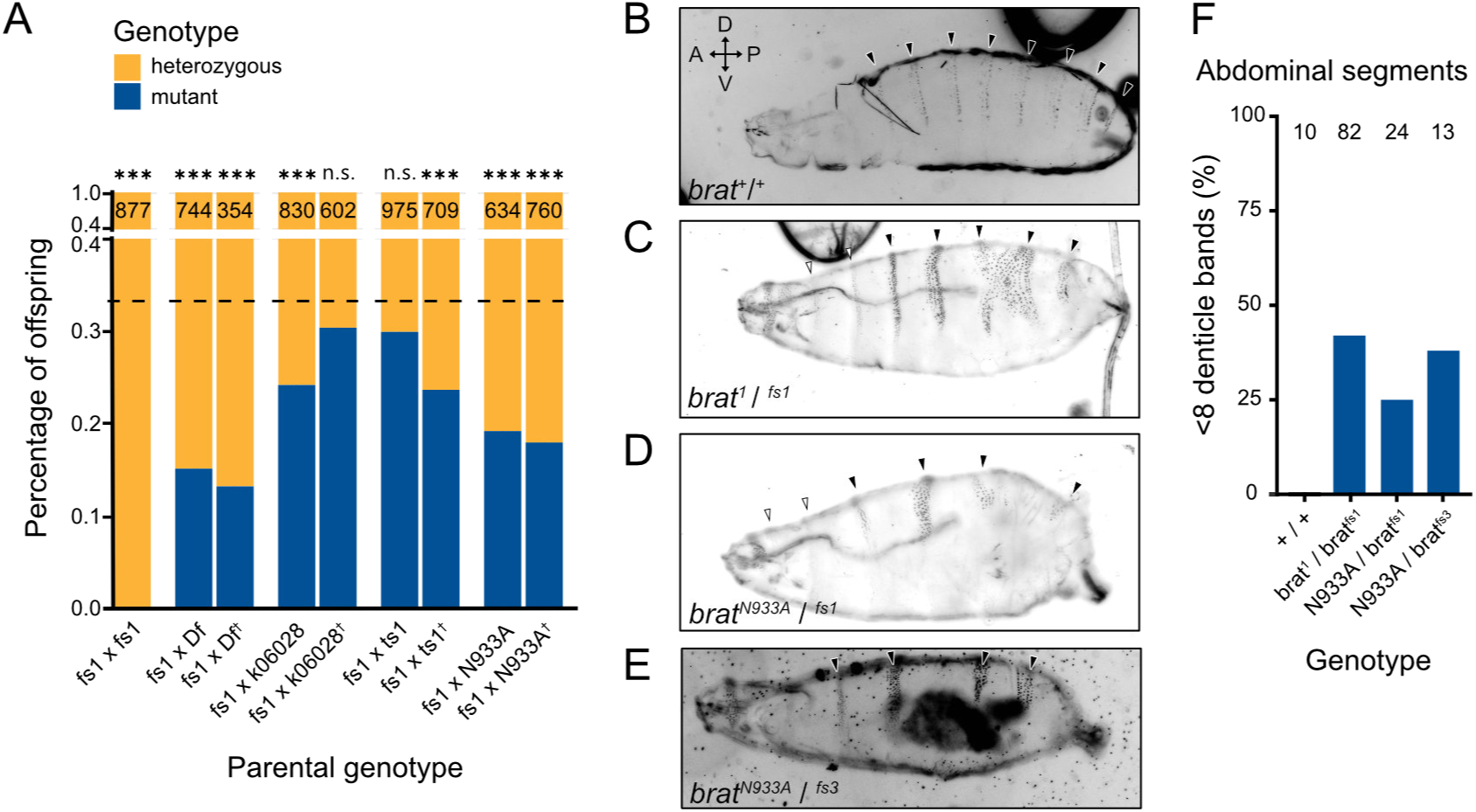
RBDmt alleles have embryonic defects. A) Stacked column graphs showing percentage of heterozygous (yellow) or mutant (blue) offspring of the indicated cross involving the known female sterile alleles. Formatting follows Figure 2. The observed fraction was compared to the expected fraction from a non-lethal combination (0.33, dashed line) by a χ^2^ Test with Bonferroni-corrected p-values for multiple comparisons (* p^adj^ < 0.01 *, < p^adj^ < 0.05, *** p^adj^ < 0.001, n.s.: not significant). B-E) Darkfield microscopy (inverted) of cuticle preparations of a representative late stage embryo. Embryos derived from mothers of the noted genotype. Anterior (A) - Posterior (P) and Ventral (V) - Dorsal (D) axis denoted. The abdominal (black arrowhead) and thoracic (white arrowhead) denticle bands are marked. F) Percentage of embryos with less than the normal number of abdominal segments. Genotypes are indicated at the bottom. Total number of embryos assessed noted at top.

Embryos from *brat^fs1^/-* mothers lack the full eight abdominal segments, marked with denticle bands, of wild-type embryos and larva (example of a hatching L1 in Figure 5(B)) ^8,9^. We confirmed this phenotype in the compound heterozygote *brat^1^/^fs1^*, which had fewer denticle bands than wild-type embryos (Figure 5(C)). This phenotype was recapitulated with the RNA-binding defective mutant allele *N933A* as a compound heterozygote of either female sterile alleles: *brat^fs1^/^N933A^* (Figure 5(D)) and *brat^fs3^/^N933A^* (Figure 5(E)). As the phenotype was less penetrant than previously reported ^8,9^, we confirmed the reported lesions of several *brat^fs1^* (G774D) and *brat^fs3^* (H802L) hemizygous larvae (Figure S4). We opted to group all embryos with less than eight denticle bands (Figure 5(F)), and in this regard the abdominal segmentation phenotype occurred at similar rates in *brat^fs1^/^N933A^* and *brat^fs3^/^N933A^* embryos as *brat^fs1^/^1^* embryos. Thus, preventing RNA-binding alone is sufficient to recapitulate the phenotype of loss-of-function *brat* alleles. These results demonstrate that the RNA-binding activity of Brat is necessary for proper embryonic development.

### RNA-binding is required to regulate Brat target mRNAs in vivo

To directly assess the role of Brat’s RNA-binding on mRNA expression levels, we measured the effect of our RBDmt alleles on the mRNA levels of natural Brat targets. We previously showed that Brat represses mRNAs encoding the vacuolar ATPase (V-ATPase) complex and glycolytic enzymes ^3,6^. Using RT-qPCR, we compared the levels of known Brat targets in total RNAs isolated from larvae. As the dosage of Brat may be crucial for RNA repression ^6^, we wished to separate and compare the levels of these targets in both mutants and heterozygotes to wild-type larvae. To unambiguously differentiate the genetically distinct heterozygotes, we used test crosses with GFP- and RFP-expressing balancer chromosomes (Figure 6(A)). Following a test cross, all genetically distinct progeny (e.g. *N933A/Df*, *Df/+*, and *N933A/+*) could be separated and RNA isolated. Additionally, all larvae were time-matched, within approximately four hours, to reduce variability simply due to developmental timing.

**Figure 6.**
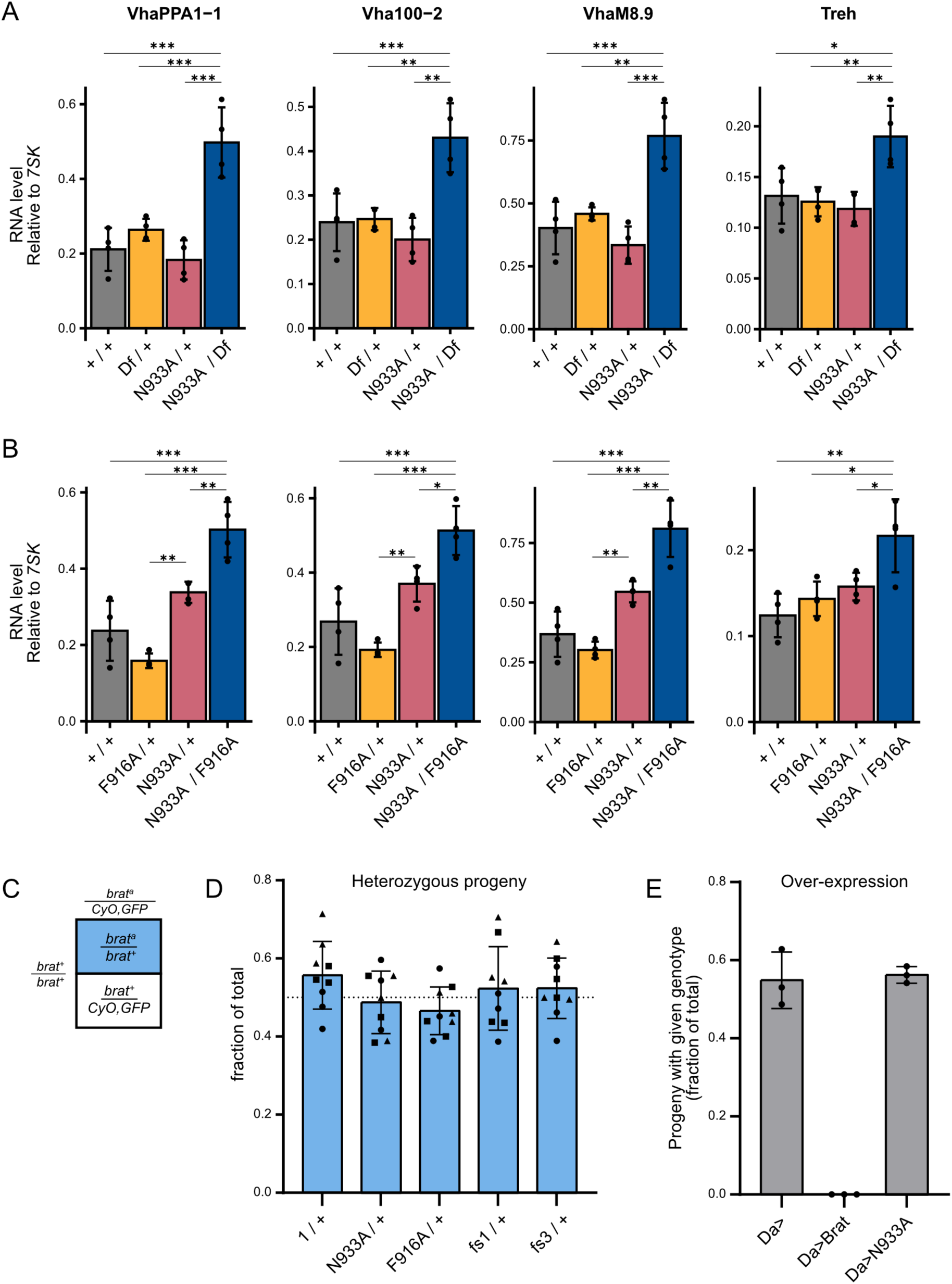
Known mRNA targets are dysregulated by RBDmt *brat* alleles. A-B) Levels of *VhaPPA1-1*, *Vha100-2*, *VhaM8.9*, and *Treh* mRNA, normalized to *7SK* ncRNA, were assessed by reverse transcription-quantitative polymerase chain reaction (RT-qPCR). Mean ± standard deviation plotted along with individual values. All viable genotypes of test crosses of *N933A* hemizygotes (A) or compound heterozygotes (B) were assessed, along with time-matched controls with wild-type alleles of *brat* (*+/+*). Significant differences in mRNA levels were determined via ANOVA with post-hoc Tukey-Kramer tests (* 0.01 < p-value < 0.05, ** 0.001 < p-value < 0.01, *** p-value < 0.001, n=4). Pairwise comparisons had not significant differences unless noted. C) Punnett square displaying the expected outcomes of a monohybrid cross of a balanced, generic brat allele (*brat^a^*) and wild-type brat alleles (*brat^+^*). In these crosses, heterozygous offspring (*brat^a^/^+^*) can be differentiated from their Cy, GFP-tagged siblings. D) Fraction of heterozygous progeny of the denoted genotype. E) Fraction of progeny with the given genotype following ubiquitous over-expression of Brat or N933A by *daughterless>Gal4* (*Da>*). The fractions of *Da>* and *Da>N933A* progeny are not significantly different (two-sample t-test, p-value 0.77). In panels E-F, the mean ± standard deviation is plotted along with individual values.

First, the levels of V-ATPase genes *VhaPPA1-1*, *Vha100-2*, and *VhaM8.9* and the metabolic gene *Treh* were compared in larva with two (*+/+*), one (*N933A / +*, *Df / +*), or zero (*N933A / Df*) wild-type copies of *brat* (Figure 5(B)). The levels of each Brat target mRNA were higher in larvae hemizygous for the RNA-binding mutant allele *brat^N933A^* (*N933A / Df*) than either heterozygote (*N933A / +*, *Df / +*) or wild-type controls (*+/+*). Furthermore, the levels of these mRNAs were not significantly different between the genetically different heterozygotes. Similarly, the levels were not different in either heterozygote compared to wild-type animals. The magnitudes of the observed increased expression levels for these target mRNAs are consistent with measurements using RNA-Seq in response to RNAi depletion of Brat in cultured cells ^6^. Furthermore, all of these targets were previously shown to be up-regulated in early embryos derived from *brat* mutant females, relative to wild-type ^3^.

To verify these mRNA differences in an additional genetic background, we also compared the mRNA levels of these genes in combinations of *brat* RBDmt alleles (Figure 5(C)). As with hemizygous animals, the levels of assessed mRNAs were higher in compound heterozygotes (*N933A / F916A*) than in larva heterozygous for RBDmt alleles (*N933A /* +, *F916A /* +) and wild-type larvae (+/+). These results demonstrate that the RNA-binding activity of Brat is necessary for proper regulation of its direct target mRNAs in vivo.

Finally, we considered the possibility that the RNA-binding defective *brat* alleles could function as a dominant negative, given their ability to interact with protein partners but not RNA. This suspicion was bolstered by the slight activation of the 8xBBS reporter observed when these alleles were ectopically expressed in cultured cells (Figure 1(F-G)). To examine this possibility, the offspring of balanced heterozygotes crossed to wild-type flies were genotyped by the presence or absence of dominant markers on the balancer chromosome (Figure 5(C)). As expected, approximately half of offspring were heterozygous, and no differences were observed between heterozygotes containing missense mutations, nonsense mutations, and the RNA-binding defective alleles (Figure 5(D), one-way ANOVA p-value = 0.1573). Furthermore, we ubiquitously over-expressed either wild-type Brat or N933A transgenes using the *daughterless-Gal4* driver (*Da>*) (Figure 5(E)). As demonstrated previously ^6^, transgenic over-expression of Brat is embryonic lethal, but flies over-expressing N933A are as viable as the driver alone (Welch’s t-test p-value 0.7778). Therefore, the data do not support RNA-binding defective Brat functioning as a dominant negative.

## Discussion

The results of this study demonstrate that the RNA-binding activity of Brat is its essential molecular function, and is necessary for embryonic and larval development, control of neural stem cell differentiation, and proper regulation of BBS-containing target mRNAs. These findings refine the conceptual framework for TRIM-NHL proteins, positioning RNA recognition as a primary and indispensable molecular function rather than an accessory activity. Since its discovery, Brat has served as a model for TRIM-NHL function in development and stem cell differentiation ^8,20–22,57,58^. Although the NHL domain was shown decades ago to be required for Brat function, its precise molecular role remained unclear ^8,25^. Early models proposed that the NHL domain functioned primarily as a protein-protein interaction platform mediating translational repression ^8,9^. This view shifted when the NHL domains of Brat and human TRIM71 were shown to bind to specific RNA sequences ^7,59^. Subsequent structural, biochemical, and transcriptome studies established sequence specific RNA recognition by NHL domains and identified transcriptome-wide Brat targets ^2–6,14,15,59,60^. Collectively, these studies provided the impetus for directly testing the biological requirement for Brat’s RNA-binding function in vivo.

Definitively assigning molecular function to a mutation requires integration of biochemical and genetic evidence. In rare instances, missense mutations can be mechanistically linked to disruption of a defined biochemical activity. For example, biochemical and structural analysis of TRIM71 provided a means of understanding how point mutations in the NHL domain of TRIM71 associated with congenital hydrocephalus disrupt RNA recognition ^61–63^. Here, precision genome editing enabled a reverse genetic strategy to mutate RNA-binding residues of Brat and test their function in the endogenous context its capacity for direct RNA recognition. RNA-binding defective *brat* alleles phenocopy strong loss-of-function and null mutations, resulting in lethality during late larval or pupal stages coincident with brain tumor formation. These phenotypes are reproducible across founders and our RNA-binding defective mutants failed to complement established *brat* mutant alleles ^20^, confirming that the defects are due to impaired Brat function. Moreover, our RNA-binding mutants recapitulated hallmark phenotypes, including neural stem cell overproliferation, embryonic segmentation defects due to *hunchback* misregulation, and increased expression of known Brat target mRNAs ^3,6^. Collectively, these findings demonstrate that Brat’s essential developmental roles depend on its capacity for direct RNA recognition.

Conceptually, our findings underscore the importance of rigorously testing assumed molecular functions. For example, although Pumilio proteins are well-established RNA-binding regulators, mutating RNA-binding residues in the *C. elegans* homolog Fbf-2 revealed that distinct phenotypes can be either RNA-binding–dependent or –independent ^64^. In the latter case, other protein partners appear to compensate in the absence of RNA-binding by Fbf-2. Given Brat’s interactions with other RNA-binding proteins, including Pumilio, Nanos, and Tis11, direct genetic testing of its RNA-binding function was necessary to distinguish primary from compensatory mechanisms ^8,34,35,65^.

Our strategy is broadly applicable to other TRIM-NHL proteins. RNA-binding has been demonstrated for *Drosophila* TRIM-NHL proteins Mei-P26 ^4,66^ and Wech ^2,4^, and *C. elegans* NCL-1 and LIN-41 ^4^. Mei-P26’s RNA-binding domain has been shown to be essential for its role in modulating the size of Type I neuroblast lineages ^67^. NCL-1 offers a unique opportunity to test the biological importance of RNA-binding in nematodes. While NCL-1 and Brat share 32.9% protein identity, its NHL domain is 83% identical, recognizes an identical RNA sequence as Brat ^2,4^, and the nucleolar phenotype of *ncl-1* mutants can be rescued with transgenic *brat* ^68^. In vertebrates, RNA-binding has been demonstrated for TRIM71 and TRIM56 ^4^. Because several TRIM-NHL family members contain RING domains and are proposed to function as E3 ubiquitin ligases (e.g. Mei-P26 ^69^ and mammalian TRIM71 ^70^), separating RNA-binding from ubiquitin ligase activity through structure-guided mutagenesis is essential for defining their primary molecular roles. Recent work supports the power of this approach in dissecting transcriptome-wide regulatory effects of TRIM71 ^71^.

Our results establish that Brat’s essential developmental functions derive from its direct regulation of target RNAs. A key next challenge is to define the full repertoire of Brat targets and delineate how repression mechanisms vary across developmental contexts. Transcriptome profiling following depletion of Brat in specific cell types has provided a necessary first step to identifying these cell-type-specific targets ^5,14,15^. Given the diverse phenotypes of our RNA-binding defective mutants, we anticipate the mutant alleles generated in this study will facilitate analysis of Brat regulatory roles in multiple cell types and developmental processes. In addition to the need to define Brat’s regulatory network, there is much we do not understand about its impact on gene expression. Future research is necessary to determine its mechanism of repression and measure its effect on mRNA decay, translation, and mRNA localization.

## Additional Information

### Author Contributions

CRediT: **Robert P. Connacher**: Conceptualization, Data curation, Formal analysis, Investigation, Visualization, Writing – original draft; **Yichao Hu**: Investigation; **Richard Roden**: Investigation; **Julia Toledo**: Investigation, Validation; **Anna DesMarais**: Investigation, Validation; **Michael O’Connor**: Resources, Writing – review & editing; **Howard D. Lipshitz**: Project administration, Writing – review & editing; **Aaron C. Goldstrohm**: Project administration, Supervision, Writing – original draft, Writing – review & editing.

### Data availability statement

The data supporting the findings of this study are available within the article [and/or] its supplementary material. Raw data and images are available upon request.

## Acknowledgements

We thank members of the Goldstrohm and O’Connor laboratories for their helpful advice and technical assistance. The anti-Brat antibodies were kindly provided by Chris Doe and Paul Lasko, and the anti-Mira antibody by Jürgen Knoblich. The anti-Elav antibody was obtained from the Developmental Studies Hybridoma Bank, University of Iowa.

## Supplementary Material

### Supplementary_Figures

Supplementary Figures S1, S2, S3, and S4.

### Additional_File_1

Oligonucleotides, plasmids, DNA fragments, RT-qPCR data & MIQE checklist, reagents, luciferase assay data & stats, fly assay data & stats, complementation assays, and genotyping of generated alleles.

## References

1. Connacher RP, Goldstrohm AC. Molecular and biological functions of TRIM-NHL RNA-binding proteins. Wiley Interdiscip Rev RNA 2021; 12:e1620.

2. Kumari P, Aeschimann F, Gaidatzis D, Keusch JJ, Ghosh P, Neagu A, Pachulska-Wieczorek K, Bujnicki JM, Gut H, Grosshans H, et al. Evolutionary plasticity of the NHL domain underlies distinct solutions to RNA recognition. Nat Commun 2018; 9:1549.

3. Laver JD, Li X, Ray D, Cook KB, Hahn NA, Nabeel-Shah S, Kekis M, Luo H, Marsolais AJ, Fung KY, et al. Brain tumor is a sequence-specific RNA-binding protein that directs maternal mRNA clearance during the Drosophila maternal-to-zygotic transition. Genome Biol 2015; 16:94.

4. Loedige I, Jakob L, Treiber T, Ray D, Stotz M, Treiber N, Hennig J, Cook KB, Morris Q, Hughes TR, et al. The Crystal Structure of the NHL Domain in Complex with RNA Reveals the Molecular Basis of Drosophila Brain-Tumor-Mediated Gene Regulation. Cell reports 2015; 13:1206–20.

5. Reichardt I, Bonnay F, Steinmann V, Loedige I, Burkard TR, Meister G, Knoblich JA. The tumor suppressor Brat controls neuronal stem cell lineages by inhibiting Deadpan and Zelda. EMBO Rep 2018; 19:102–17.

6. Connacher RP, Roden RT, Huang K-L, Korte AJ, Yeruva S, Dittbenner N, DesMarais AJ, Weidmann CA, Randall TA, Williams J, et al. The TRIM-NHL RNA-binding protein Brain Tumor coordinately regulates expression of the glycolytic pathway and vacuolar ATPase complex. Nucleic Acids Res 2024; 52:12669–88.

7. Loedige I, Stotz M, Qamar S, Kramer K, Hennig J, Schubert T, Loffler P, Langst G, Merkl R, Urlaub H, et al. The NHL domain of BRAT is an RNA-binding domain that directly contacts the hunchback mRNA for regulation. Genes Dev 2014; 28:749–64.

8. Sonoda J, Wharton RP. Drosophila Brain Tumor is a translational repressor. Genes Dev 2001; 15:762–73.

9. Cho PF, Gamberi C, Cho-Park YA, Cho-Park IB, Lasko P, Sonenberg N. Cap-dependent translational inhibition establishes two opposing morphogen gradients in Drosophila embryos. Curr Biol 2006; 16:2035–41.

10. Eichhorn SW, Subtelny AO, Kronja I, Kwasnieski JC, Orr-Weaver TL, Bartel DP. mRNA poly(A)-tail changes specified by deadenylation broadly reshape translation in Drosophila oocytes and early embryos. Elife 2016; 5:e16955.

11. Subtelny AO, Eichhorn SW, Chen GR, Sive H, Bartel DP. Poly(A)-tail profiling reveals an embryonic switch in translational control. Nature 2014; 508:66–71.

12. Vastenhouw NL, Cao WX, Lipshitz HD. The maternal-to-zygotic transition revisited. Development 2019; 146.

13. Zamore PD, Williamson JR, Lehmann R. The Pumilio protein binds RNA through a conserved domain that defines a new class of RNA-binding proteins. Rna 1997; 3:1421–33.

14. Landskron L, Steinmann V, Bonnay F, Burkard TR, Steinmann J, Reichardt I, Harzer H, Laurenson A-S, Reichert H, Knoblich JA. The asymmetrically segregating lncRNA cherub is required for transforming stem cells into malignant cells. Elife 2018; 7:e31347.

15. Bonnay F, Veloso A, Steinmann V, Köcher T, Abdusselamoglu MD, Bajaj S, Rivelles E, Landskron L, Esterbauer H, Zinzen RP, et al. Oxidative Metabolism Drives Immortalization of Neural Stem Cells during Tumorigenesis. Cell 2020; 182:1490–1507.e19.

16. Homem CC, Knoblich JA. Drosophila neuroblasts: a model for stem cell biology. Development 2012; 139:4297–310.

17. Janssens DH, Lee CY. It takes two to tango, a dance between the cells of origin and cancer stem cells in the Drosophila larval brain. Semin Cell Dev Biol 2014; 28:63–9.

18. Bello B, Reichert H, Hirth F. The brain tumor gene negatively regulates neural progenitor cell proliferation in the larval central brain of Drosophila. Development 2006; 133:2639–48.

19. Beaucher M, Goodliffe J, Hersperger E, Trunova S, Frydman H, Shearn A. Drosophila brain tumor metastases express both neuronal and glial cell type markers. Dev Biol 2007; 301:287–97.

20. Lee CY, Wilkinson BD, Siegrist SE, Wharton RP, Doe CQ. Brat is a Miranda cargo protein that promotes neuronal differentiation and inhibits neuroblast self-renewal. Dev Cell 2006; 10:441–9.

21. Betschinger J, Mechtler K, Knoblich JA. Asymmetric segregation of the tumor suppressor brat regulates self-renewal in Drosophila neural stem cells. Cell 2006; 124:1241–53.

22. Bowman SK, Rolland V, Betschinger J, Kinsey KA, Emery G, Knoblich JA. The tumor suppressors Brat and Numb regulate transit-amplifying neuroblast lineages in Drosophila. Dev Cell 2008; 14:535–46.

23. Wright TR, Hodgetts RB, Sherald AF. The genetics of dopa decarboxylase in Drosophila melanogaster. I. Isolation and characterization of deficiencies that delete the dopa-decarboxylase-dosage-sensitive region and the alpha-methyl-dopa-hypersensitive locus. Genetics 1976; 84:267–85.

24. Wright TR, Bewley GC, Sherald AF. The genetics of dopa decarboxylase in Drosophila melanogaster. II. Isolation and characterization of dopa-decarboxylase-deficient mutants and their relationship to the alpha-methyl-dopa-hypersensitive mutants. Genetics 1976; 84:287–310.

25. Arama E, Dickman D, Kimchie Z, Shearn A, Lev Z. Mutations in the beta-propeller domain of the Drosophila brain tumor (brat) protein induce neoplasm in the larval brain. Oncogene 2000; 19:3706–16.

26. McCrady E, Tolin DJ. Effects of Ddc cluster lethal alleles on ovary growth, attachment, and egg production in Drosophila. J Exp Zool 1994; 268:469–76.

27. Schupbach T, Wieschaus E. Female sterile mutations on the second chromosome of Drosophila melanogaster. II. Mutations blocking oogenesis or altering egg morphology. Genetics 1991; 129:1119–36.

28. Spradling AC, Stern D, Beaton A, Rhem EJ, Laverty T, Mozden N, Misra S, Rubin GM. The Berkeley Drosophila Genome Project gene disruption project: Single P-element insertions mutating 25% of vital Drosophila genes. Genetics 1999; 153:135–77.

29. Stathakis DG, Pentz ES, Freeman ME, Kullman J, Hankins GR, Pearlson NJ, Wright TR. The genetic and molecular organization of the Dopa decarboxylase gene cluster of Drosophila melanogaster. Genetics 1995; 141:629–55.

30. Wright TR, Beermann W, Marsh JL, Bishop CP, Steward R, Black BC, Tomsett AD, Wright EY. The genetics of dopa decarboxylase in Drosophila melanogaster. IV. The genetics and cytology of the 37B10-37D1 region. Chromosoma 1981; 83:45–58.

31. Kurzik-Dumke U, Phannavong B, Gundacker D, Gateff E. Genetic, cytogenetic and developmental analysis of the Drosophila melanogaster tumor suppressor gene lethal(2)tumorous imaginal discs (1(2)tid). Differentiation 1992; 51:91–104.

32. Woodhouse E, Hersperger E, Shearn A. Growth, metastasis, and invasiveness of Drosophila tumors caused by mutations in specific tumor suppressor genes. Dev Genes Evol 1998; 207:542–50.

33. Komori H, Xiao Q, McCartney BM, Lee CY. Brain tumor specifies intermediate progenitor cell identity by attenuating beta-catenin/Armadillo activity. Development 2014; 141:51–62.

34. Komori H, Golden KL, Kobayashi T, Kageyama R, Lee CY. Multilayered gene control drives timely exit from the stem cell state in uncommitted progenitors during Drosophila asymmetric neural stem cell division. Genes Dev 2018; 32:1550–61.

35. Newton FG, Harris RE, Sutcliffe C, Ashe HL. Coordinate post-transcriptional repression of Dpp-dependent transcription factors attenuates signal range during development. Development 2015; 142:3362–73.

36. Komori H, Rastogi G, Bugay JP, Luo H, Lin S, Angers S, Smibert CA, Lipshitz HD, Lee C-Y. mRNA decay pre-complex assembly drives timely cell-state transitions during differentiation. Cell Rep 2025; 44:115138.

37. Gratz SJ, Ukken FP, Rubinstein CD, Thiede G, Donohue LK, Cummings AM, O’Connor-Giles KM. Highly specific and efficient CRISPR/Cas9-catalyzed homology-directed repair in Drosophila. Genetics 2014; 196:961–71.

38. DeLuca SZ, Spradling AC. Efficient Expression of Genes in the Drosophila Germline Using a UAS Promoter Free of Interference by Hsp70 piRNAs. Genetics 2018; 209:381–7.

39. flyCRISPR – CRISPR Information and resources for Drosophila researchers [Internet]. [cited 2023 Nov 7]; Available from: https://flycrispr.org/

40. Huang W, Massouras A, Inoue Y, Peiffer J, Ràmia M, Tarone AM, Turlapati L, Zichner T, Zhu D, Lyman RF, et al. Natural variation in genome architecture among 205 Drosophila melanogaster Genetic Reference Panel lines. Genome Res 2014; 24:1193–208.

41. Mackay TFC, Richards S, Stone EA, Barbadilla A, Ayroles JF, Zhu D, Casillas S, Han Y, Magwire MM, Cridland JM, et al. The Drosophila melanogaster Genetic Reference Panel. Nature 2012; 482:173–8.

42. Morgan TH, Sturtevant AH, Muller HJ, Bridges CB. The mechanism of Mendelian heredity [Internet]. Revised Edition. New York: Henry Holt and Company; 1923. Available from: http://resource.nlm.nih.gov/05321170R

43. Alexandre C. Cuticle preparation of Drosophila embryos and larvae. Methods Mol Biol 2008; 420:197–205.

44. Hoyer’s Medium. Cold Spring Harb Protoc 2011; 2011:pdb.rec12429.

45. Bustin SA, Benes V, Garson JA, Hellemans J, Huggett J, Kubista M, Mueller R, Nolan T, Pfaffl MW, Shipley GL, et al. The MIQE guidelines: minimum information for publication of quantitative real-time PCR experiments. Clin Chem 2009; 55:611–22.

46. Arias AM. Drosophila melanogaster and the Development of Biology in the 20th Century [Internet]. In: Dahmann C, editor. Drosophila: Methods and Protocols. Totowa, NJ: Humana Press; 2008 [cited 2026 Feb 19]. page 1–25. Available from: 10.1007/978-1-59745-583-1_1

47. Arbeille E, Bashaw GJ. Brain Tumor promotes axon growth across the midline through interactions with the microtubule stabilizing protein Apc2. PLoS Genet 2018; 14:e1007314.

48. Horváth B, Kalinka AT. Effects of larval crowding on quantitative variation for development time and viability in Drosophila melanogaster. Ecol Evol 2016; 6:8460–73.

49. Loop T, Leemans R, Stiefel U, Hermida L, Egger B, Xie F, Primig M, Certa U, Fischbach KF, Reichert H, et al. Transcriptional signature of an adult brain tumor in Drosophila. BMC Genomics 2004; 5:24.

50. Boone JQ, Doe CQ. Identification of Drosophila type II neuroblast lineages containing transit amplifying ganglion mother cells. Dev Neurobiol 2008; 68:1185–95.

51. Bier E, Vaessin H, Younger-Shepherd S, Jan LY, Jan YN. deadpan, an essential pan-neural gene in Drosophila, encodes a helix-loop-helix protein similar to the hairy gene product. Genes Dev 1992; 6:2137–51.

52. Ceron J, González C, Tejedor FJ. Patterns of cell division and expression of asymmetric cell fate determinants in postembryonic neuroblast lineages of Drosophila. Dev Biol 2001; 230:125–38.

53. Ikeshima-Kataoka H, Skeath JB, Nabeshima Y, Doe CQ, Matsuzaki F. Miranda directs Prospero to a daughter cell during Drosophila asymmetric divisions. Nature 1997; 390:625–9.

54. Shen CP, Jan LY, Jan YN. Miranda is required for the asymmetric localization of Prospero during mitosis in Drosophila. Cell 1997; 90:449–58.

55. Robinow S, White K. Characterization and spatial distribution of the ELAV protein during Drosophila melanogaster development. J Neurobiol 1991; 22:443–61.

56. Arvola RM, Weidmann CA, Tanaka Hall TM, Goldstrohm AC. Combinatorial control of messenger RNAs by Pumilio, Nanos and Brain Tumor Proteins. RNA Biol 2017; :1–12.

57. Neumuller RA, Betschinger J, Fischer A, Bushati N, Poernbacher I, Mechtler K, Cohen SM, Knoblich JA. Mei-P26 regulates microRNAs and cell growth in the Drosophila ovarian stem cell lineage. Nature 2008; 454:241–5.

58. Xiao Q, Komori H, Lee CY. klumpfuss distinguishes stem cells from progenitor cells during asymmetric neuroblast division. Development 2012; 139:2670–80.

59. Loedige I, Gaidatzis D, Sack R, Meister G, Filipowicz W. The mammalian TRIM-NHL protein TRIM71/LIN-41 is a repressor of mRNA function. Nucleic Acids Res 2013; 41:518–32.

60. Wang, Y., Yu, Z., Wang, M., Liu, C. P., Yang, N., Xu, R. M. 5EX7: Crystal structure of Brat NHL domain in complex with an 8-nt hunchback mRNA. 2015;

61. Furey CG, Choi J, Jin SC, Zeng X, Timberlake AT, Nelson-Williams C, Mansuri MS, Lu Q, Duran D, Panchagnula S, et al. De Novo Mutation in Genes Regulating Neural Stem Cell Fate in Human Congenital Hydrocephalus. Neuron 2018; 99:302–314 e4.

62. Liu Q, Novak MK, Pepin RM, Maschhoff KR, Worner K, Chen X, Zhang S, Hu W. A congenital hydrocephalus-causing mutation in Trim71 induces stem cell defects via inhibiting Lsd1 mRNA translation. EMBO Rep 2023; 24:e55843.

63. Duy PQ, Furey CG, Kahle KT. Trim71/lin-41 Links an Ancient miRNA Pathway to Human Congenital Hydrocephalus. Trends Mol Med 2019; 25:467–9.

64. Carrick BH, Crittenden SL, Linsley M, Dos Santos SJC, Wickens M, Kimble J. The PUF RNA-binding protein, FBF-2, maintains stem cells without binding to RNA. RNA 2025; 31:623–32.

65. Harris RE, Pargett M, Sutcliffe C, Umulis D, Ashe HL. Brat promotes stem cell differentiation via control of a bistable switch that restricts BMP signaling. Dev Cell 2011; 20:72–83.

66. Salerno-Kochan A, Horn A, Ghosh P, Nithin C, Kościelniak A, Meindl A, Strauss D, Krutyhołowa R, Rossbach O, Bujnicki JM, et al. Molecular insights into RNA recognition and gene regulation by the TRIM-NHL protein Mei-P26. Life Sci Alliance 2022; 5:e202201418.

67. Hu Y, Yang X, Lipshitz HD. The TRIM-NHL RNA-binding protein MEI-P26 modulates the size of Drosophila Type I neuroblast lineages. Genetics 2025; 229:iyaf015.

68. Frank DJ, Edgar BA, Roth MB. The Drosophila melanogaster gene brain tumor negatively regulates cell growth and ribosomal RNA synthesis. Development 2002; 129:399–407.

69. Ferreira A, Boulan L, Perez L, Milan M. Mei-P26 mediates tissue-specific responses to the Brat tumor suppressor and the dMyc proto-oncogene in Drosophila. Genetics 2014; 198:249–58.

70. Rybak A, Fuchs H, Hadian K, Smirnova L, Wulczyn EA, Michel G, Nitsch R, Krappmann D, Wulczyn FG. The let-7 target gene mouse lin-41 is a stem cell specific E3 ubiquitin ligase for the miRNA pathway protein Ago2. Nat Cell Biol 2009; 11:1411–20.

71. Welte T, Tuck AC, Papasaikas P, Carl SH, Flemr M, Knuckles P, Rankova A, Buhler M, Grosshans H. The RNA hairpin binder TRIM71 modulates alternative splicing by repressing MBNL1. Genes Dev 2019; 33:1221–35.

