## Supplementary Figures for "RNA-binding is the essential biological function of the *Drosophila* protein Brat"

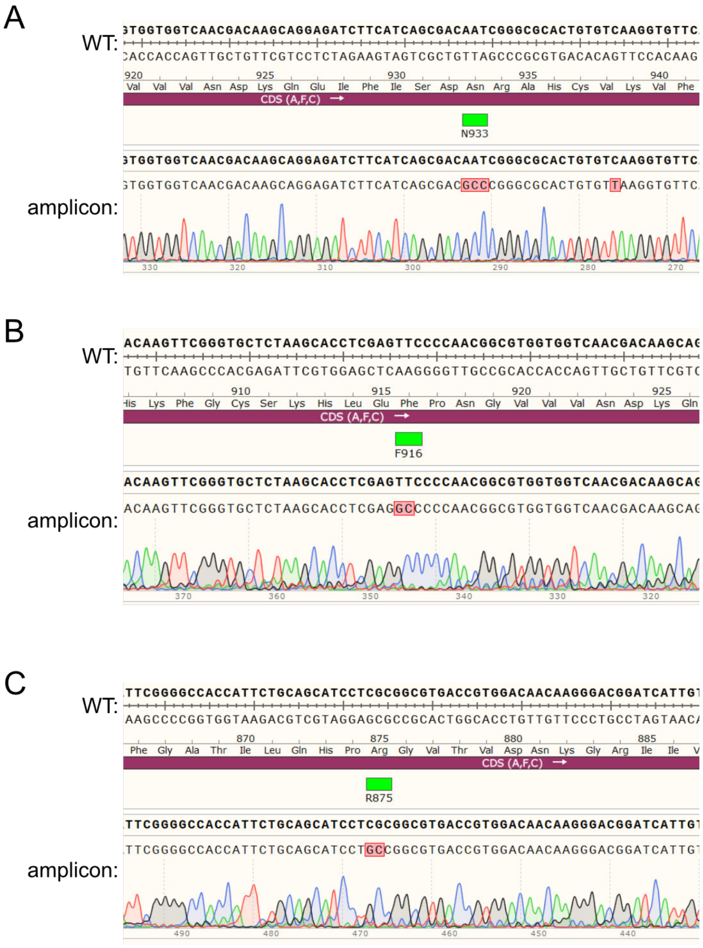


**Figure S1. The RBDmt *brat* alleles were confirmed by Sanger sequencing**
Chromatograms from select Sanger Sequencing reactions highlighting the RBDmt alleles. The NHL region was amplified from genomic DNA of A) *N933A*, B) *F916A*, and C) *R875A* hemi- and homozygotes as described in Figure 1. Note the synonymous mutation in V938 in panel A is from the Scarless Gene Editing strategy.

A

| allele | Description | Source | Nature of allele Identified |
|---|---|---|---|
| $brat^{1}$ | Q872* (PTC) | Wright (1976) | Connacher (2024) |
| $brat^{18}$ | Q507* (PTC) | Wright (1976) | This publication (Kyoto stock #101379) |
| *k06028* | *lacW* P-element insertion in 5′UTR | Spradling (1999) | Arama (2000) |
| $brat^{fs1}$ | G774D (missense) | Wright (1976) | Sonoda & Wharton (2001) |
| $brat^{fs3}$ | H802L (missense) | Schubach & Wieschaus (1991) | Arama (2000), GenBank AF195873.1 |
| $brat^{ts1}$ | G860D (missense) | Wright (1976) | Arama (2000), GenBank AF195872 |

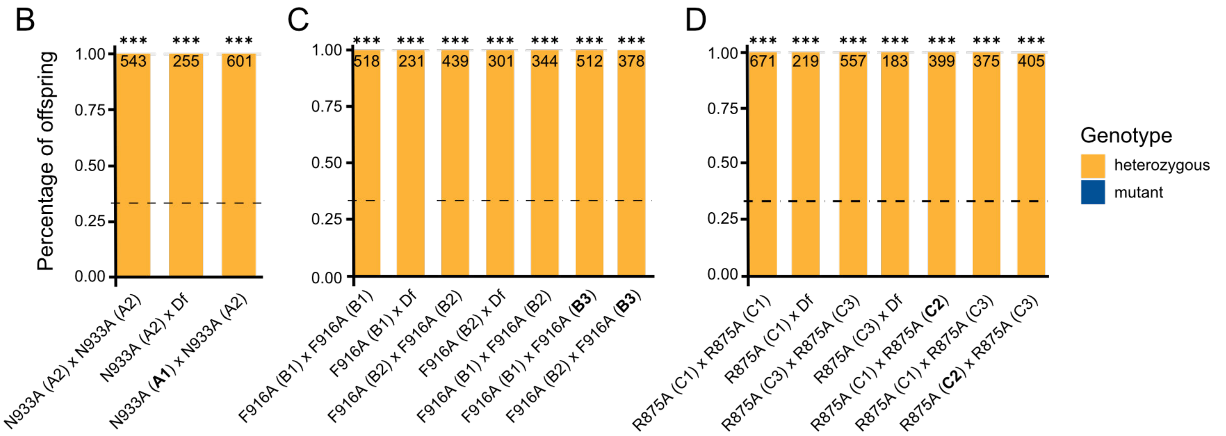


**Figure S2. Additional complementation assays**
A) Table of *brat* alleles and descriptions used in this study. Alleles were classified as premature truncation codons (PTC), missense, or transgenic insertions. The source of the allele references the attribution from flybase.org. B-D) Stacked column graphs showing percentage of heterozygous (yellow) or mutant (blue) offspring of the indicated cross, involving additional isogenic founders of *N933A* (B), *F916A* (C), and *R875A* (D) alleles. Formatting follows Figure 2. Bold: The isogenic lines N933A (A1), F916A (B3), and R875A (C2) are used to represent the RBDmt alleles throughout the manuscript. The observed fraction was compared to the expected fraction from a non-lethal combination (0.33, dashed line) by a $\chi^2$ Test with Bonferroni-corrected p-values for multiple comparisons (*** $p^{adj} < 0.001$).

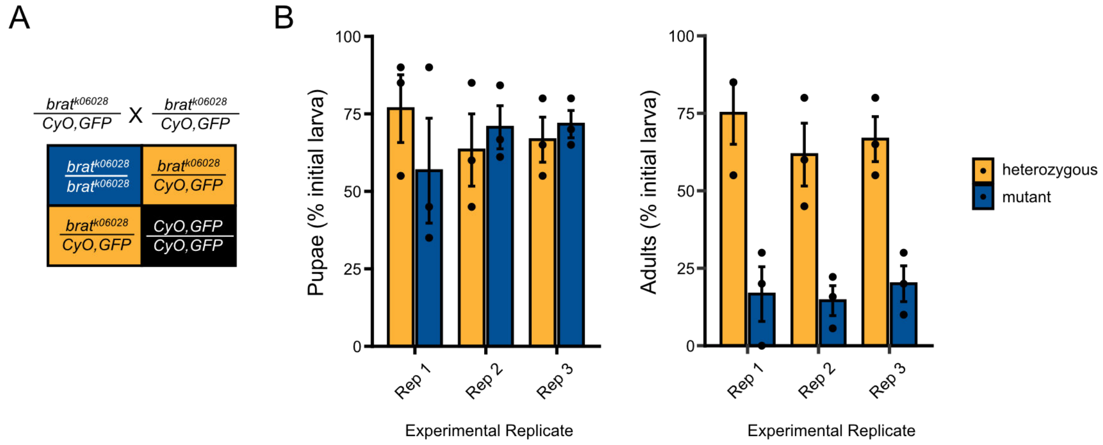


**Figure S3. Reproducibility of lethal phase analysis assays**

A) Punnett square displaying the expected outcomes of a monohybrid cross of the *brat*$^{k06028}$ allele using balancer chromosomes, following Figure 2A. B) Rates of larvae that survive to pupal stages (left) and adulthood (right), as a percentage of initial larva. The class of genotype is noted. Each experimental replicate (cross) included three technical replicates (vials) of sorted GFP$^{+}$ or GFP$^{-}$ first instar larvae.

A

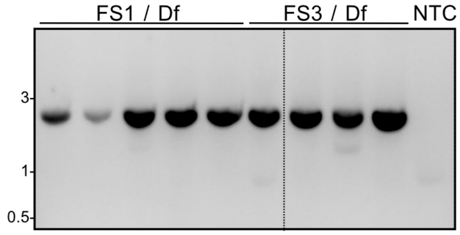


B

| mutation | genomic coordinates | FS1 / Df larvae | FS3 / Df larvae |
|---|---|---|---|
| G774[GGC]>D774[GAC] | 2L:19170466(+)G>A | 5/5 | 0/4 |
| R761[CGA]>R761[CGG] | 2L:19170428(+)A>G | 5/5 | 4/4 |
| F815[TTT]>F815[TTC] | 2L:19170167(+)T>C | 5/5 | 0/4 |
| H802[CAT]>L802[CTT] | 2L:19170550(+)A>T | 0/5 | 4/4 |
| G643[GGC]>S643[AGC] | 2L:19170072(+)G>A | 0/5 | 4/4 |
| T669[ACG]>T669[ACT] | 2L:19170152(+)G>T | 0/5 | 4/4 |
| R761[CGA]>R761[CGG] | 2L:19170428(+)A>G | 0/5 | 4/4 |

**Figure S4. The female sterile alleles were confirmed by Sanger sequencing**
A) Agarose gel electrophoresis of PCR products amplified from genomic DNA of individual $brat^{fs1}$ (FS1 / Df) or $brat^{fs3}$ (FS3 / Df) larvae. No amplification was observed in the no-template control (NTC). Sizes are in kilobases. B) Mutations (and corresponding genomic coordinates) identified by Sanger Sequencing of these PCR products. The number of larva with these lesions, out of the total tested, is shown.
